# The CoREST complex inhibitor, corin, leads to decreased tumor growth, increased cellular differentiation and extended lifespan in atypical teratoid rhabdoid tumor xenograft models

**DOI:** 10.1101/2024.12.14.628381

**Authors:** Anupa Geethadevi, Nikhil Vaidya, Robert J. Fisher, Tyler Findlay, Yiming Deng, Khoa Pham, Vikas Kumar, Samuel Beck, Calixto-Hope G. Lucas, Charles G. Eberhart, Jinchong Xu, Philip A. Cole, Jeffrey Rubens, Marianne Collard, Eric H. Raabe, Rhoda M. Alani

**Author notes:** To whom correspondence should be addressed: Marianne Collard, PhD, 609 Albany Street, J-504, Boston, Massachusetts 02118,; Eric H. Raabe, MD, PhD, Division of Pediatric Oncology, Johns Hopkins University School of Medicine, 1800 Orleans St., 11th Floor, Baltimore, MD 21287,; Rhoda M. Alani, MD, 609 Albany Street, J-507, Boston, Massachusetts 02118.

## Abstract

**Background:** Atypical teratoid rhabdoid tumor (ATRT) is the most common malignant brain tumor in infants, and more than 60% of children with ATRT die from their tumor. ATRT is associated with mutational inactivation/deletion of *SMARCB1*, a member of the SWI/SNF chromatin remodeling complex, suggesting that epigenetic events play a critical role in tumor development and progression. Moreover, disruption of SWI/SNF allows unopposed activity of epigenetic repressors, which contribute to tumorigenicity. We therefore explored the role of the CoREST repressor complex in ATRT.

**Methods:** We evaluated the effects of the bifunctional LSD1/HDAC1/2 small molecule CoREST inhibitor, corin, on ATRT tumor cell growth, apoptosis, differentiation, gene expression and chromatin accessibility.

**Results:** We found that corin inhibited the growth of ATRT cells regardless of their epigenetic subgroup, and was associated with increased tumor cell apoptosis and differentiation. ATAC-seq showed increases in chromatin accessibility in corin-treated ATRT cells, with changes seen at genes associated with neuronal differentiation and synaptic function. RNA-seq confirmed increased expression of neuronal differentiation genes and decreased DNA replication/cell cycle-associated genes in ATRT cells treated with corin. Corin suppressed orthotopic ATRT tumor growth, leading to significant extension of lifespan. In addition, increased histone acetylation (H3K9ac, H3K27ac) and methylation (H3K4Me1) was seen in corin-treated ATRT orthotopic xenografts, consistent with on-target pharmacodynamics.

**Conclusion:** The CoREST inhibitor, corin, suppresses tumor growth, induces differentiation, and promotes apoptosis in ATRT, leading to significantly increased survival of mice bearing ATRT orthotopic xenografts. Our results suggest a potential application of CoREST complex inhibitors in patients with ATRT.

**Key Points:**

- CoREST complex inhibition by corin leads to decreased cell growth and increased apoptosis in ATRT
- Corin promotes chromatin accessibility and neuronal differentiation in ATRT
- Corin inhibits tumor growth and extends lifespan in ATRT animal models

**Importance of the Study:** Loss of function of SMARCB1 is a hallmark of ATRT which leads to dysfunction of the mammalian SWI/SNF complex and an inability to counteract epigenetic repressor complexes. The CoREST complex functions as a chromatin remodeling complex that represses neuronal differentiation genes during development. Inhibition of the CoREST complex by corin in ATRT leads to decreased tumor cell growth, induction of apoptosis and increased survival of mice bearing ATRT orthotopic xenografts. These changes are associated with increased chromatin accessibility and expression of genes associated with neuronal differentiation. Corin therefore reverses the primary block of differentiation that maintains a stem cell state in ATRT, which contributes to tumorigenesis. These studies significantly improve our understanding of how to therapeutically address the underlying epigenetic drivers of ATRT and support further development of corin for ATRT.

## Introduction

Atypical teratoid rhabdoid tumor (ATRT) is the most common malignant brain tumor in infants. ATRT has a very poor prognosis with a median survival of around 17 months and a 5-year survival rate of 32%^1,2^. Current therapeutic interventions include surgical removal, intensive chemotherapy, stem cell transplants and radiation, which lead to lifelong toxicity for survivors^1,2^. Although ATRT is a highly aggressive malignancy, the only recurrent genetic event is bi-allelic loss-of-function mutation in *SMARCB1* (95% of cases) or *SMARCA4* (5% of cases) ^3–5^ which encode subunits of the mammalian SWI/SNF chromatin remodeling complex.^6^

The SWI/SNF complex regulates chromatin compaction and DNA accessibility via alteration of nucleosome positioning and occupancy in an ATP-dependent manner and is disrupted in over 20% of human cancers.^7^ SMARCB1 is a critical mediator of SWI/SNF-mediated activation of gene enhancer and promotor activity.^8^ Chromatin regulation is critical for the maintenance of timely and appropriate gene expression, and epigenetic regulators play key roles in normal development and oncogenesis.^9,10^ While the SWI/SNF complex facilitates DNA accessibility, histone-modifying enzymes post-translationally alter the N-terminal tails of histones with consequent alterations in chromatin structure and binding of transcriptional regulatory proteins. During normal neural development, the Polycomb repressive complex (PRC2) and SWI/SNF complex create a balance between maintenance of a stem cell state and neuronal differentiation.^11^ Absence of SMARCB1 compromises the full activity of the SWI/SNF complex leading to an array of epigenetic derangements in the neurodevelopmental process that alters the physiological balance from differentiation towards maintenance of a stem cell state. A hallmark of ATRT is high-level expression of stem cell factors such as LIN28A, LIN28B, HMGA2 and MYC, emphasizing the imbalance between stemness and differentiation in this tumor.^12,13^

Recent studies have determined that despite the absence of SMARCB1 in ATRT, the residual SWI/SNF activity opposes EZH2 except at loci that are occupied by a strong transcriptional repressor, REST (RE1 Silencing Transcription factor). REST is closely associated with neuronal stem cells and is upregulated in ATRT,^14^ and its activity is thought to be mediated by association with the CoREST repressor complex.^15–17^ RCOR1/RCOR2/RCOR3 are scaffold proteins that recruit one copy of the LSD1 (lysine-specific demethylase 1) and one copy of histone deacetylases HDAC1 or HDAC2 and this tripartite organization comprises the CoREST complex, a neuro-developmentally important chromatin repressor complex, to site-specific domains on chromatin leading to transcriptional repression.^18,19^

Corin is a potent and selective bifunctional inhibitor of the CoREST repressor complex reported in 2018.^20^ Corin inhibits tumor cell growth in melanoma, cutaneous squamous cell carcinoma, colon cancer, diffuse intrinsic pontine glioma (DIPG), malignant peripheral nerve sheath tumor (MPNST) and breast cancer through its dual ability to target both the histone deacetylase (HDAC 1,2) and lysine-specific demethylase 1 (LSD1) active sites of the CoREST complex, increasing its selectivity for this HDAC complex.^20–25^ Corin has increased potency and an improved therapeutic index versus the corresponding class I HDAC inhibitor (entinostat/ MS-275) alone or in combination with LSD1 inhibitor (GSK-LSD1) in some cancer types.^20,23^ Additionally, corin inhibits tumor cell plasticity and reverses therapy resistance in human melanoma cells.^23^ We therefore hypothesized that corin would restore epigenetic balance in ATRT by reactivating silenced differentiation programs, leading to growth arrest.

We find that corin reprogrammed the epigenetic landscape in ATRT leading to inhibition of tumor cell growth, induction of apoptosis and neuronal differentiation with potent in vivo effects in promoting survival of mice bearing tumor xenografts. Our data suggest that CoREST complex inhibitors may prove useful in the treatment of this aggressive malignancy.

## Materials and Methods

### Cell Culture and Reagents

The ATRT cell lines CHLA-02-ATRT, CHLA-04-ATRT, CHLA-05-ATRT, and CHLA-06-ATRT were a gift from Anat Erdreich-Epstein.^26^ These cell lines are now available at ATCC (Manassas, VA) for purchase. The ATRT cell line, BT37, was derived from a serially passaged xenograft derived from a patient with ATRT.^12,27^ BT12-ATRT and CHLA266-ATRT cell line were procured from Children’s Oncology Group cell repository (Lubbock, TX). The culture conditions of each cell line are given in Supplementary Table 1. All cell lines were authenticated periodically by INI1 western blotting (showing loss of expression in all cases) and by STR DNA testing. The details of antibodies and primers used in the experiments are mentioned in Supplementary Tables 2, 3, and 4.

### Compounds

The HDAC inhibitor entinostat/MS-275 (Cat# HY-12163) and corin (Cat#: HY-111048) were purchased from MedChemExpress (Monmouth Junction, NJ, USA). Both were prepared as previously described.^20^ All stock solutions were dissolved in DMSO. The concentration of corin used for each cell line and time point of various experiments is shown in Supplementary Table 5.

### Cell growth inhibition assay

For determining the potency and efficacy of corin, ATRT cells were plated at a density of 10,000 cells per well in a 96-well plate and serial concentrations of corin were added in each well. Cell Counting Kit-8 (CCK-8) (CK04, Dojindo) was used according to the manufacturer’s suggested protocol. After 72 h of corin treatment, 10 µL of CCK-8 solution was added to each well and the plates were incubated at 37 °C in 5% CO_2_ for 60 min. Absorbance at 450 nm was measured as indicative of cell viability. Corin IC_50_ values for each cell line were determined based on the decrease in the total number of viable cell count, using Graph Pad Prism v10.0. Cells treated with DMSO were used as vehicle control. The relative viability and percent growth inhibition was determined using a flow-cytometry based method after 72 hours of corin treatment (MUSE Count & Viability Kit, Cat# MCH100102, Cytek Biosciences Inc., Fremont, CA) as per the manufacturer’s protocol (see Supplementary Methods). Growth inhibition assay for entinostat/MS-275 (HDAC1/3 inhibitor) was also simultaneously performed for the ATRT cell lines to estimate the comparative efficacy of Corin with MS275 using the CCK-8 Assay Kit as described previously.

### MTS growth assays

For the MTS assays, cells were seeded in a 96-well plate at a density of 2,000 cells per well for BT37 and CHLA06, and 5,000 cells per well for CHLA05 and treated with corin or DMSO. MTS reagent (Cat# G3581, Promega Corp., Madison, WI, USA) was added at various time points (0 h, 24 h, 48 h and 96 h/ 120 h), incubated for 4 h and the absorbance was measured at 590 nm. The absorbances of all time points were normalized to that at 0 h to estimate the relative growth of ATRT cells for over 3-4 days, with and without corin.

### Annexin V/ Dead cell assay flow cytometry

Cells were plated at a density of 200,000 cells per well in a 6-well plate with or without corin. The apoptotic cells in each group were measured using MUSE cell analyzer (MUSE Annexin V & Dead Cell Kit, Cat# MCH100105, Cytek Biosciences Inc., Fremont, CA) according to the manufacturer’s instructions (see Supplementary Methods).

### ATAC-Seq and analysis

100,000 cells were seeded and treated with corin or DMSO. DNA was extracted, tagmented, purified, and adapter ligated using the ATAC-seq Kit (ActiveMotif #53150) according to the manufacturer’s instructions. Sequencing was done by Azenta Life Sciences (MA, USA), and reads were trimmed using the fastp command in fastp v.0.23.4. The detailed protocol for both sequencing and analysis is given in Supplementary Methods.

### RNA-seq and Analysis

RNA was extracted from ATRT cells, BT37 and CHLA06 treated with corin or DMSO (RNAeasy Mini kit, Cat# 73404, Qiagen, Hilden, Germany) using the protocol described in Supplementary Methods Section. RNA samples passing the initial QC parameters (A260/280 and A230/280 ≥ 2.0, RIN score > 7), were selected for RNA sequencing using the Illumina NovoSeq 6000 platform (Novogene Corporation Inc, CA, USA). Analysis of RNA sequencing data is described in detail in the Supplementary Methods.

### ChIP-qPCR

BT37 and CHLA06 cells were treated with corin or DMSO for 24 hours in T175 flasks. ChIP was conducted as previously described.^28^ Briefly, cells were fixed with 1% formaldehyde (for H3K9ac IP) or crosslink gold and 1% formaldehyde (for RCOR1 and LSD1 IPs). Chromatin was extracted and sheared to approximately 200 bp using the Diagenode Bioruptor Pico (Diagenode, Denville, NJ, USA) using 20 cycles (crosslink gold-fixed samples) or 10 cycles (formaldehyde-fixed samples) according to manufacturer’s instructions (1 cycle = 30 seconds on/30 seconds off). Approximately 800-2000 µg protein, calculated from the sonicated chromatin, was used per immunoprecipitation reaction with antibodies listed in Supplementary Table 4. Immunoprecipitated complexes were washed and DNA was eluted, treated with Proteinase K, and reverse crosslinked. ChIP DNA was purified using a PCR Purification Kit (Qiagen, Germantown, MD, USA). qPCR was performed as described above with primers listed in Supplementary Table 4 and results were analyzed using the 1% input method. Two biological replicates were performed for each cell line.

### Immunofluorescence assays

Immunofluorescence staining for BrdU and Proliferating cell nuclear antigen (PCNA) was done to assess cell proliferation and Cleaved caspase-3 (CC3) was done to detect apoptosis. Cells were treated with corin or DMSO for 72 h. For BrdU immunofluorescence, cells were incubated in 10 nM BrdU (Cat# B23151, Thermo Fisher Scientific, Waltham, MA, USA) for 4 h, collected and fixed in cytospin fluid. Fixed cells were then cytospun onto positively charged glass slides, permeabilized in 0.1% TritonX in PBS, incubated in 1N HCl, blocked using 5% normal goat serum and incubated in anti-BrdU antibody overnight. This was followed by incubation in Cy3-labeled secondary antibody, counterstaining in DAPI (Cat# 10236276001, Roche Life Sciences, Sigma-Aldrich, MI, USA) and mounting using Prolong Gold Anti-fade mounting media (Cat# P36930, Thermo Fisher Scientific, Waltham, MA, USA). For CC3 and PCNA, cells were similarly processed, except that the DNA denaturation step with 1N HCl was omitted. Images were captured using NIS Elements F5.21.00 software attached to Nikon Motic AE31 microscope. Images were blinded to the treatment condition and the number of immuno-positive cells were counted using the ImageJ software, and normalized to the total number of cells as visualized using DAPI counterstaining.

### Western immunoblotting

ATRT cells were collected after 72 h of treatment with corin or DMSO, and lysed using lysis buffer containing RIPA buffer, protease and phosphatase inhibitors. Protein quantification was done using the Bradford method (Cat #5000006, Bio-Rad Laboratories Inc., Hercules, CA, USA). Western blotting was performed as previously described.^29^ The detailed protocol is given in Supplementary Methods.

### Neuronal differentiation

CHLA06 cells were treated with corin for 72 h and Wheat Germ Agglutinin (WGA) staining was done which helped define the morphogenic differentiation observed in these cells. Cells were fixed in 4% paraformaldehyde, washed in PBS and stained with 5 µg/ml WGA (ThermoFisher # W11261) for 10 min, and then counterstained with DAPI. For the immunofluorescence of structural (MAP2, Beta-3-tubulin/TUBB3) and functional neuronal markers (Synaptophysin/ SYP), CHLA06 and BT37 were treated with corin in chamber slides while CHLA05 was treated in T25 flasks for these experiments. After 72 h of corin treatment, immunofluorescence was performed as previously described^30^.

### Intracranial xenograft tumors

For animal care and anesthesia, “Principles of laboratory animal care” (NIH publication No. 86-23, revised 1985) were followed, using a protocol approved by the Johns Hopkins Animal Care and Use Committee, in compliance with the United States Animal Welfare Act regulations and Public Health Service Policy. Intracranial xenografts of ATRT cell lines BT37 and CHLA06 were established in anesthetized animals using the guide-screw system as previously described.^31^ Two types of experiments were performed using these intracranial xenograft models: Short-term experiments to check for target engagement of the drug, and long-term survival experiments to assess the survival advantage rendered by corin in the tumor-bearing mice. 3 µl of corin (0.03 mg corin per mouse) or DMSO (vehicle) was administered once (short-term) or weekly (long-term survival) through the guide screw. The experimental details are described in the Supplementary Methods section.

### Histology, Immunohistochemistry and Immunofluorescence of FFPA mice xenografts

Xenograft tumors were embedded in paraffin and processed for hematoxylin and eosin (H&E) staining and for Ki67 immunohistochemistry by the Johns Hopkins Histopathology Core. The extent of tumor infiltration and the number of mitotic bodies were quantified by a pathologist who was blinded to the groups. In each case with viable tumor present, the cross-sectional area of the largest tumor focus on a 10x microscopic field was quantified in pixel^2^ using the polygon tool in QuPath software (v0.3.2).^32^ We performed immunofluorescence staining on the FFPE sections of orthotopic xenografts to detect histone modifications, neuronal differentiation and apoptosis after corin treatment. The sections were deparaffinized in xylene followed by decreasing grades of ethanol, and finally in water to hydrate the sections. Steam-mediated antigen retrieval was performed on these sections using 0.1 M citrate buffer for 20 min. The sections were allowed to cool down, dipped in PBS, followed by permeabilization in 0.1% Triton-X /PBS. Sections were then blocked in 5% NGS, incubated with primary antibody overnight, followed by secondary antibody and counterstained with DAPI nuclear stain. The sections were then mounted in Prolong Gold Anti-fade mounting media. The details of all the antibodies used are given in Supplementary Table 2 and 3.

### Kaplan Meier Survival Analysis

RNA expression data from the Pediatric Brain Tumor Atlas (PBTA, Provisional) dataset was accessed from PedcBioPortal (https://pedcbioportal.org/). Data was filtered for patients with ATRT (CANCER_TYPE = Atypical Teratoid Rhabdoid Tumor) and overall survival (OS_MONTHS) data was extracted along with expression data for genes of interest. Patients were separated into tertiles (n=20/group) by gene expression level, with corresponding “high” or “low” gene expression groups. Kaplan Meier curves were plotted with GraphPad Prism v10 and analyzed by the Log-rank (Mantel-Cox) statistical test.

### Statistical Analysis

All results from the in vitro experiments are reported as Mean and standard deviation, and groups were compared using Student t-test or Mann-Whitney test, depending on the group distribution. A p-value < 0.05 was considered a statistical difference among the compared groups. Kaplan-Meier survival analysis followed by Log-rank (Mantel-Cox) test was performed to compare the median survival among the corin- and vehicle-treated groups of mice bearing orthotopic ATRT xenografts. All data analysis was performed using Graph Pad prism v 10.1.

## Results

### The CoREST inhibitor, corin, blocks tumor cell growth and promotes apoptosis in ATRT

Previous studies suggested that REST-associated transcriptional repression may be a significant mediator of tumorigenesis in SMARCB1-null tumors like ATRT,^14^ and REST transcription may be mediated by the CoREST repressor complex.^15–17^ We sought to determine the effect of our bi-functional HDAC1-LSD1 inhibitor, corin, on ATRT proliferation versus the HDACi, entinostat/MS-275, using a panel of seven established ATRT cell lines (Figure 1A – C). These cell lines represent the spectrum of ATRT molecular subgroups including ATRT-MYC, ATRT-SHH, and ATRT-TYR (Figure 1C).^33^ Corin significantly inhibited the growth of all cell lines, with variability in IC_50_s between cell lines (0.201-5.18μM) (Figure 1A).The IC_50_s for corin were consistently lower (2.22- 7.52-fold) than those of entinostat/MS-275 in all ATRT cell lines evaluated except for BT37 where the IC_50_s for MS-275 and corin were nearly identical (Figure 1B, 1C). A flow-cytometry based cell viability method showed a similar dose-dependent reduction in viability with corin (Supplementary Figure 1A), and the concentration of corin required to inhibit cell proliferation was lower than that required to reduce cell viability or cause cell death (Supplementary Figure 1B). Corin led to decreased proliferation as measured by BrdU incorporation in ATRT cells (corin vs DMSO - BT37: −11.2-fold, p = 0.0007; CHLA05: −2.8 fold, p = 0.001; CHLA06: 3.5 fold, p < 0.0001) (Figure 1D), and ATRT cells treated with corin showed sustained decrease in growth over a period of 96 – 120 hours as seen in MTS growth assays (Supplementary Figure 1C). Corin-treated ATRT cells also showed decreased expression of proliferating cell and nuclear antigen (PCNA), a marker of DNA replication and cell proliferation (corin vs DMSO- BT37: −1.8 fold, p < 0.0001; CHLA05: 1.9 fold, p = 0.008) (Supplementary Figure 1D).

**Figure 1.**
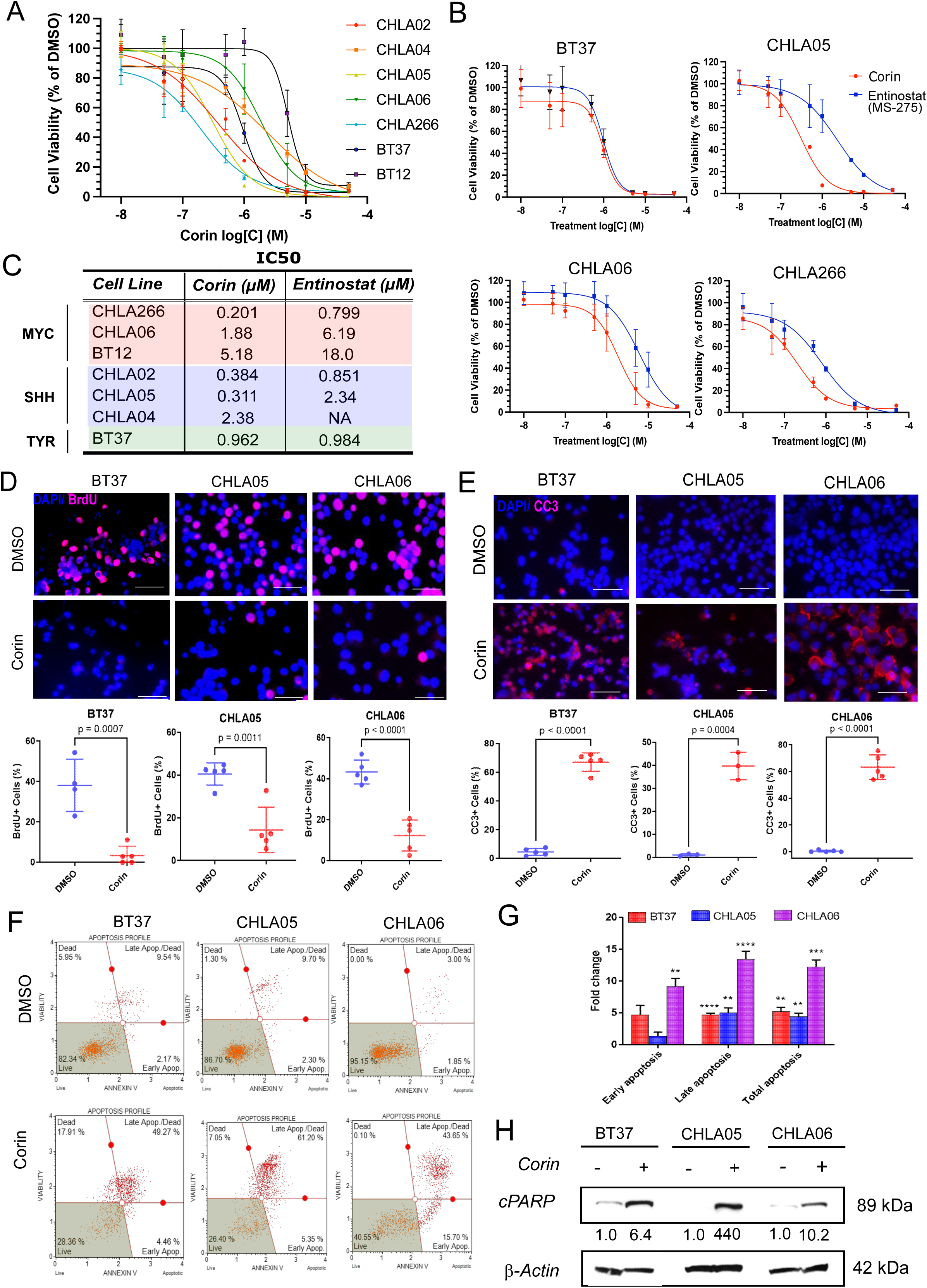
Corin effectively targets ATRT cells *in vitro*. **(A)** Dose-response curves for ATRT cell lines treated with corin for 72 h. **(B)** Relative cell viability of ATRT cell lines treated with corin versus the HDAC inhibitor, entinostat/ MS-275. **(C)** Table showing comparative IC_50_s of corin and entinostat in ATRT cell lines sorted by molecular subtype. IC_50_ was calculated based on the total viable cell count relative to DMSO in ATRT cell lines. **(D)** BrdU immunostaining and quantification of ATRT cells treated with DMSO versus corin. Scale 50 µm. **(E)** Cleaved caspase 3 (CC3) immunostaining for apoptosis and quantification of ATRT cells treated with DMSO versus corin. Scale 50 µm. **(F)** MUSE Annexin V apoptosis assay of ATRT cells treated with DMSO or corin. **(G)** Quantification of Annexin V apoptosis assay for corin treatment of ATRT cells in (F). Each bar represents mean of fold change of number of cells in various stages of apoptosis following corin treatment of ATRT cells versus DMSO in each cell line. Asterisks (*) above each bar indicates the statistical significance of corin treated cells compared with DMSO. **(H)** Western blot of cPARP expression in ATRT cells following 72 h treatment with DMSO versus corin. The band density normalized to beta actin is shown below each band. Statistical Analysis: unpaired, two-tailed student’s *t* test. All data represents mean ± SD. ** p ≤ 0.01; *** p ≤ 0.001; **** p ≤ 0.0001.

Cell lines were further evaluated for apoptosis following corin treatment using CC3 immunostaining (Figure 1E), Annexin V staining (Figure 1F, G) and cleaved PARP (cPARP) expression (Figure 1H). Corin significantly induced apoptosis in all cell lines tested following 72 hours of treatment (Figure 1E – 1G) with associated increases in cPARP (Figure 1H). Corin did not induce apoptosis in brain organoids derived from induced pluripotent stem cells even at 5 μM concentration, as demonstrated by lack of cPARP expression after 72 h (Supplementary Figure 1E).

### Selective inhibition of HDAC1/ LSD1 reprograms the epigenetic landscape and enhances chromatin accessibility in ATRT

We have previously shown that inhibiting the CoREST complex led to increases in histone H3 lysine-9 acetylation (H3K9ac), histone H3 lysine-27 acetylation (H3K27ac) and histone H3 lysine-4 methylation (H3K4me) in melanoma cell lines^23^ and therefore sought to determine whether corin treatment of ATRT cells would alter H3K9/H3K27 acetylation, methylation and chromatin accessibility. Five ATRT cell lines were treated with corin for 24 hours and H3K9ac, H3K27ac, H3K4Me1 and H3K27Me3 levels were evaluated by Western blot (Figure 2A). Interestingly, while expected increases in H3K4Me1, H3K9ac and H3K27ac were seen in all cell lines, increases in H3K27Me3 were also noted across all cell lines (Figure 2A). To evaluate chromatin accessibility changes associated with corin treatment of ATRT, BT37 and CHLA06 cells treated with DMSO or corin for 24 hours were evaluated by ATAC-seq (Figure 2B – D). As expected, corin treatment of both ATRT cell lines led to an increase in chromatin accessibility (Figure 2B). Annotation of the accessibility peaks across the genome revealed large gains in chromatin accessibility within gene bodies following corin treatment of both cell lines (Figure 2C) with significantly increased chromatin accessibility following corin treatment noted in BT37 cells (Figure 2D). HOMER motif analysis of ATAC-seq peaks gained with corin treatment for 24 h in BT37 and CHLA06 ATRT cell lines identified enriched motifs associated with zinc-finger transcription factors including CTCF/CTCFL, KLF6 and KLF7 (Figure 2E). Gene ontology analysis of genes with gains in ATAC-seq peaks within their gene bodies +/- 3Kbp of the genes in BT37 (Figure 2F) and CHLA06 (Figure 2G) cells treated with corin for 24 hours identified peaks associated with neuronal differentiation and extracellular matrix reorganization.

**Figure 2.**
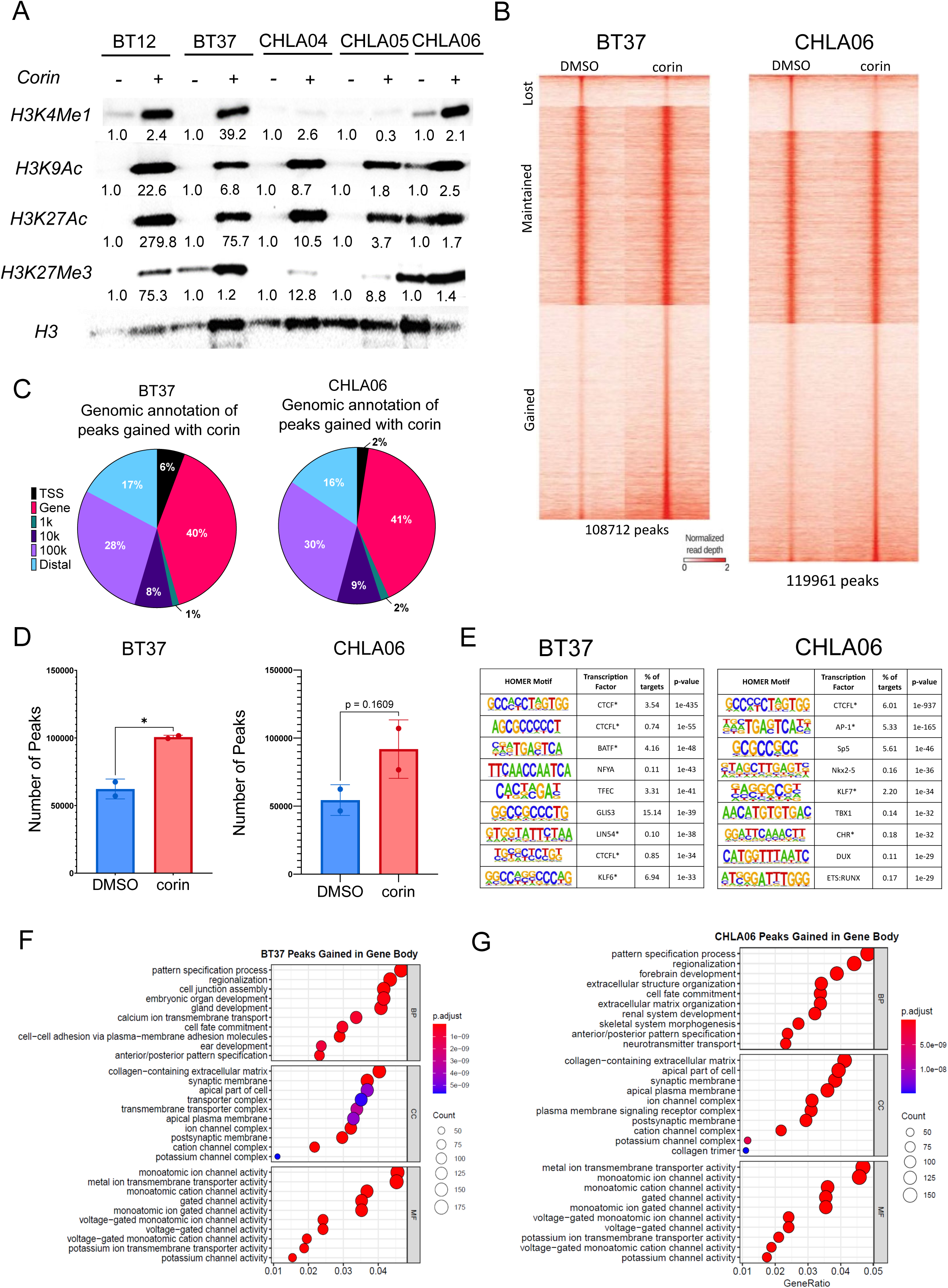
Corin increases chromatin accessibility in ATRT cells. **(A)** Western blot of H3K4Me1, H3K9Ac, H3K27Ac and H3K27Me3, with H3 as loading control, in ATRT cells treated with DMSO or corin for 24 h. **(B)** Tornado plots of ATAC-seq signals lost, maintained, or gained with corin treatment for 24 h compared to DMSO in BT37 and CHLA06 cells. **(C)** Genomic annotations of peaks gained with corin treatment, and **(D)** quantification of total peaks. **(E)** HOMER motif analysis results of ATAC-seq peaks gained with corin treatment for 24 h compared to DMSO in BT37 or CHLA06 cells. Asterisks indicate motif families common between the top-10 listed motifs for BT37 and CHLA06 cells. **(F-G)** Gene ontology analysis of genes with gains in ATAC-seq peaks within their gene bodies +/- 3Kbp of the genes in **(F)** BT37 or **(G)** CHLA06 cells treated with corin for 24 h compared to DMSO. Asterisks indicate the top displayed gene sets common between BT37 and CHLA06 cells. Statistical Analysis: unpaired, two-tailed student’s *t* test. All data represents mean ± SD. \**P*<0.05. (n=2).

### Corin treatment of ATRT cells promotes neuronal differentiation

We next sought to determine the global transcriptional effects of CoREST inhibition in BT37 and CHLA06 ATRT cell lines to try to discern the functional consequences of CoREST inhibition in ATRT. Corin treatment resulted in morphological changes in ATRT cell lines in culture, such as elongation of cellular processes suggesting neuronal differentiation, which was further confirmed by staining the cell membrane of CHLA06 cells using Alexa 488 wheat germ agglutinin (WGA) (Figure 3A). To further evaluate the effect of corin, we treated ATRT cells with corin for 24 hours and evaluated by RNA- seq.

**Figure 3.**
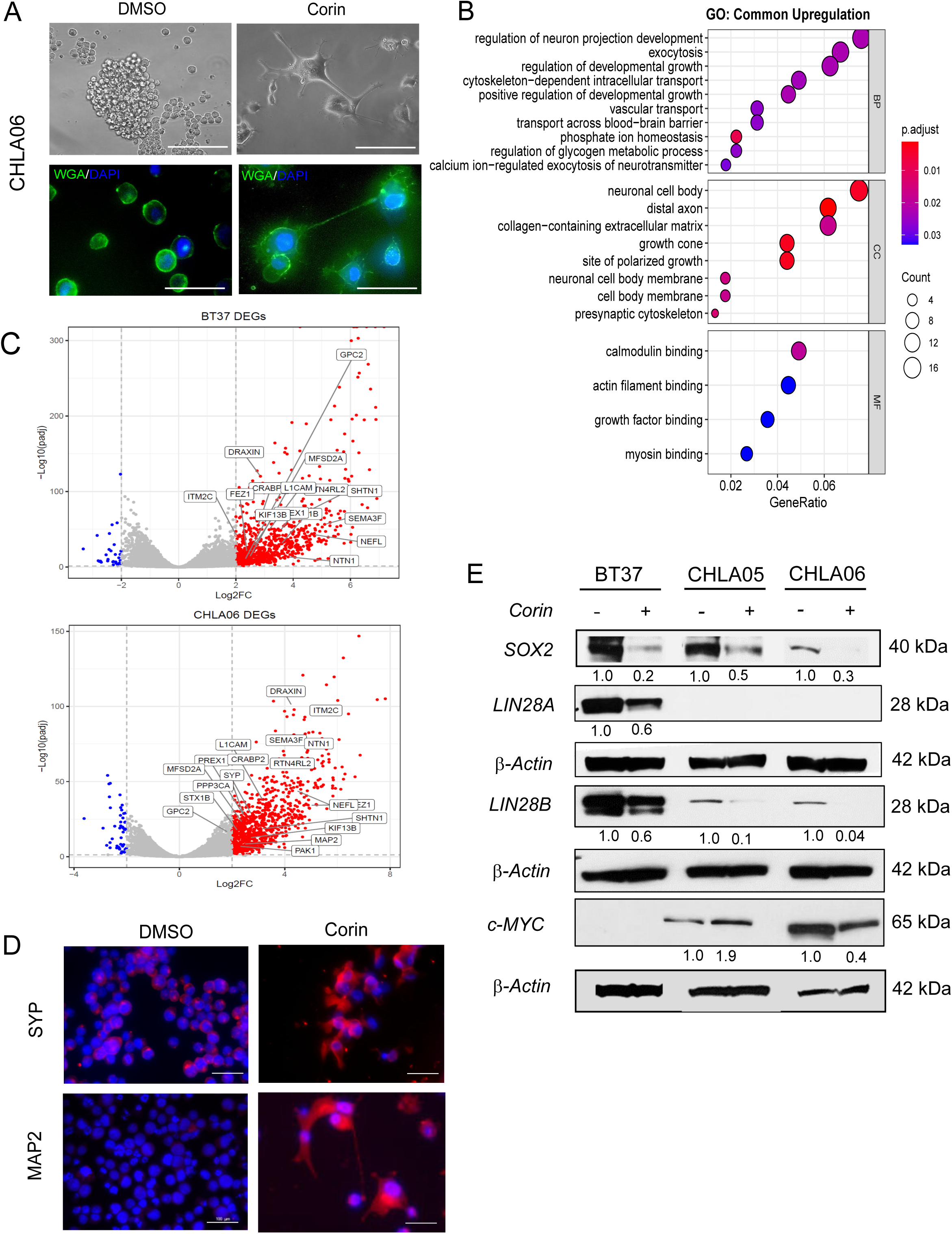
Corin induces neuronal differentiation and decreases stemness in ATRT cells. **(A)** Wheat germ agglutinin staining, counterstained with DAPI, of CHLA06 cells treated with DMSO versus corin for 72 h. Scale 50 µm. **(B)** Pathway analysis on whole transcriptome sequencing of ATRT cells treated with corin for 24 h reveals upregulation of neurogenesis and neuron developmental pathways (PFDR=0.05). **(C)** Volcano plot illustrating important neuronal differentiation genes upregulated by corin in BT37 and CHLA06 cells. **(D)** Immunofluorescence staining of neuronal differentiation markers, Synaptophysin (SYP), and MAP2, following 72 h of DMSO or corin treatment in CHLA06 cells. Scale 50 µm. **(E)** Western blot of stemness markers, SOX2, LIN28A, LIN28B and c-MYC, in ATRT cells following 72 h of DMSO or corin treatment.

Consistent with inhibition of the repressive functions of CoREST^18^ we found that most genes altered in expression by corin were upregulated (Supplementary Figure 2A). Among the 1554 genes upregulated (P FC cut-off p < 0.05, Log2FC ≥ 2) in both BT37 and CHLA06 cells, 261 genes were commonly upregulated (Supplementary Figure 3B). Gene set enrichment analysis (GSEA) of differentially expressed genes revealed a significant positive enrichment (p < 0.001) for SMARCB1 targets that are repressed in ATRT as defined by Erkek *et al*. These gene promoters are associated with a significant enrichment for neuronal differentiation transcription factor motifs and suggest that corin treatment upregulates a gene signature traditionally repressed in SMARCB1-deficient tumors. Gene Ontology (GO) analysis of common upregulated pathways in these ATRT cell lines following corin treatment revealed the highest common upregulated biological process was *regulation of neuron projection development.* The top upregulated cellular processes were *neuronal cell body* and *distal axon* (Figure 3B) with *genes associated with neuronal differentiation* being the most significantly increased in expression after corin treatment (Figure 3C). Gene set enrichment analysis (GSEA) for genes associated with neuronal projection development demonstrated significant upregulation in both BT37 and CHLA06 cells treated with corin (Supplementary Figure 2D).

Immunofluorescence of neuronal differentiation markers Synaptophysin (SYP) and Microtubule-associated Protein 2 (MAP2) increased in corin-treated cells compared to control, which corroborated the RNA-seq data (Figure 3D). Although the ATRT-SHH subgroup cell line CHLA05 did not demonstrate morphological differentiation following corin treatment, we observed significantly higher expression of neuronal differentiation markers (Supplementary Figure 2E). Additionally, the structural protein, Beta-3-tubulin, which is almost exclusively present in neurons, was also highly expressed in ATRT cells treated with corin (Supplementary Figure 2F).

Overexpression of stem cell factors such as LIN28A, LIN28B, SOX2 and MYC are a hallmark of ATRT, and downregulation of stem cell factors leads to improved survival in mouse models.^12,13,29^ We found that 72 hour treatment with corin downregulated the stem cell factors LIN28B and SOX2 in all ATRT cell lines, and LIN28A in BT37 (Figure 3E). Corin treatment also led to dramatically reduced MYC protein expression in CHLA06, which is an ATRT-MYC subgroup cell line (Figure 3E).

### Corin promotes ATRT differentiation as evidenced by increased neuronal projection development genes, *NEFL* and *L1CAM*

GO-based enrichment analysis of the Neuron Projection Development pathway further revealed critical genes associated with neuronal differentiation that were significantly upregulated in corin-treated BT37 and CHLA06 cells (Figure 4A). Among these neuronal differentiation-associated genes, we identified Neurofilament Light polypeptide (*NEFL/ NFL*) which is involved in regulating axon growth and branching, and Neural cell adhesion molecule L1 (*L1CAM/ NCAML1*) which is involved in mediating neuronal migration and inter-neuronal interactions, to be upregulated in corin-treated ATRT cells compared to control. We confirmed increased L1CAM and NEFL in corin treated ATRT by Western blot (Figure 4B). Integration of RNA-seq and ATAC-seq signal tracks revealed that *NEFL* upregulation is associated with an increase in promoter chromatin accessibility in both BT37 and CHLA06 cell lines (Figure 4C). To assess if this effect was direct, we designed *NEFL* promoter-specific primers and conducted ChIP-qPCR using antibodies against H3K9ac and CoREST components, LSD1 and RCOR1, following corin treatment. We found that corin significantly increased H3K9ac marks in both cell lines and identified an enrichment for CoREST occupancy at this region (Figure 4D and 4E). Additionally, we observed a similar pattern at the *L1CAM* promoter (Figure 4F-H) suggesting that CoREST complex plays a critical role in directly regulating the expression of these gene targets. To validate the importance of neural differentiation in ATRT, we evaluated patient survival using the PedcBioPortal dataset. Kaplan-Meier survival analysis found that patients whose tumors expressed high levels of *L1CAM* had improved survival compared to those with low *L1CAM* (p-value = 0.0025 by Log-rank test) (Figure 4I). Patients with ATRT tumors that expressed high levels of *NEFL* mRNA had a median survival of 43 months compared to 23 months for patients with low *NEFL* (Figure 4J).

**Figure 4.**
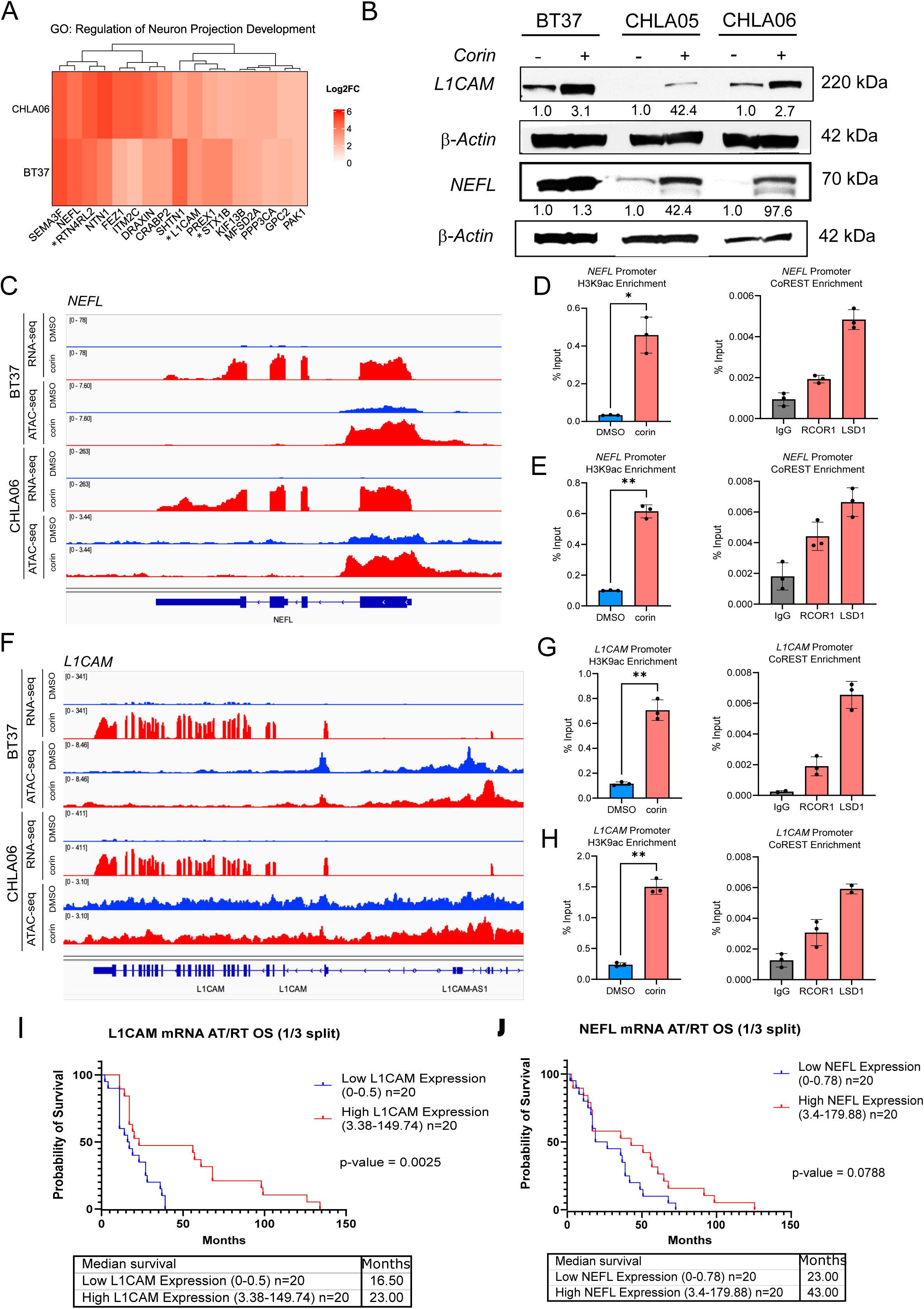
**(A)** GO-based enrichment analysis of the Neuron Projection Development pathway reveals critical genes involved in mediating neuronal differentiation in corin-treated ATRT cells, BT37 and CHLA06. **(B)** Western immunoblotting showing increased expression of neuronal differentiation markers, L1CAM and NEFL, after corin treatment. **(C)** Signal tracks of RNA-seq (RPKM) and ATAC-seq (FKPM) signal peaks at the *NEFL* locus in BT37 and CHLA06 cells treated with DMSO or corin for 24 h. **(D)** H3K9ac occupancy at the *NEFL* promotor in BT37 cells treated with DMSO or corin for 24 h (left) with occupancy of RCOR1 or LSD1 compared to IgG in BT37 control cells at the same locus (right). **(E)** H3K9ac occupancy at the *NEFL* promotor in CHLA06 cells treated with DMSO or corin for 24 h (left) with occupancy of RCOR1 or LSD1 compared to IgG in CHLA06 control cells at the same locus (right). **(F)** Signal tracks of RNA-seq (RPKM) and ATAC-seq (FKPM) signal peaks at the *L1CAM* locus in BT37 and CHLA06 cells treated with DMSO or corin for 24 h. **(G)** H3K9ac occupancy at the *L1CAM* promotor in BT37 cells treated with DMSO or corin for 24 h (left) with occupancy of RCOR1 or LSD1 compared to IgG in BT37 control cells at the same locus (right). **(H)** H3K9ac occupancy at the *L1CAM* promotor in CHLA06 cells treated with DMSO or corin for 24 h (left) with occupancy of RCOR1 or LSD1 compared to IgG in CHLA06 control cells at the same locus (right). Kaplan-Meier survival curves showing the survival advantage of increased **(I)** *L1CAM*, and **(J)** *NEFL* expression in primary ATRT patients using data derived from Pediatric Brain Tumor Network dataset. (*p < 0.05, **p < 0.01).

### Short-term treatment with corin leads to decreased tumor cell growth, increased apoptosis and cellular differentiation in ATRT orthotopic xenograft models

To determine whether corin treatment increases apoptosis and engages with the histone targets in vivo, we performed a short-term study of corin activity in ATRT orthotopic xenografts in athymic nude mice. ATRT cells were injected through a guide-screw implanted in the right cerebral hemisphere and corin was injected through the guide screw after tumors were well established (Figure 5A). Tumors treated with corin for 24 hours demonstrated decreased numbers of mitotic figures (Figure 5B), increased CC3 (Figure 5C) and increased cPARP expression (Figure 5E) indicative of decreased proliferation and increased apoptosis. Furthermore, tumors demonstrated increased expression of H3K9Ac (Figure 5D), H3K4Me1 and H3K27Ac (Figure 5E) illustrating that corin engaged with its targets in vivo. Tumors treated with corin also demonstrated induction of neuronal differentiation as indicated by expression of the mature neuronal marker synaptophysin (Figure 5F).

**Figure 5.**
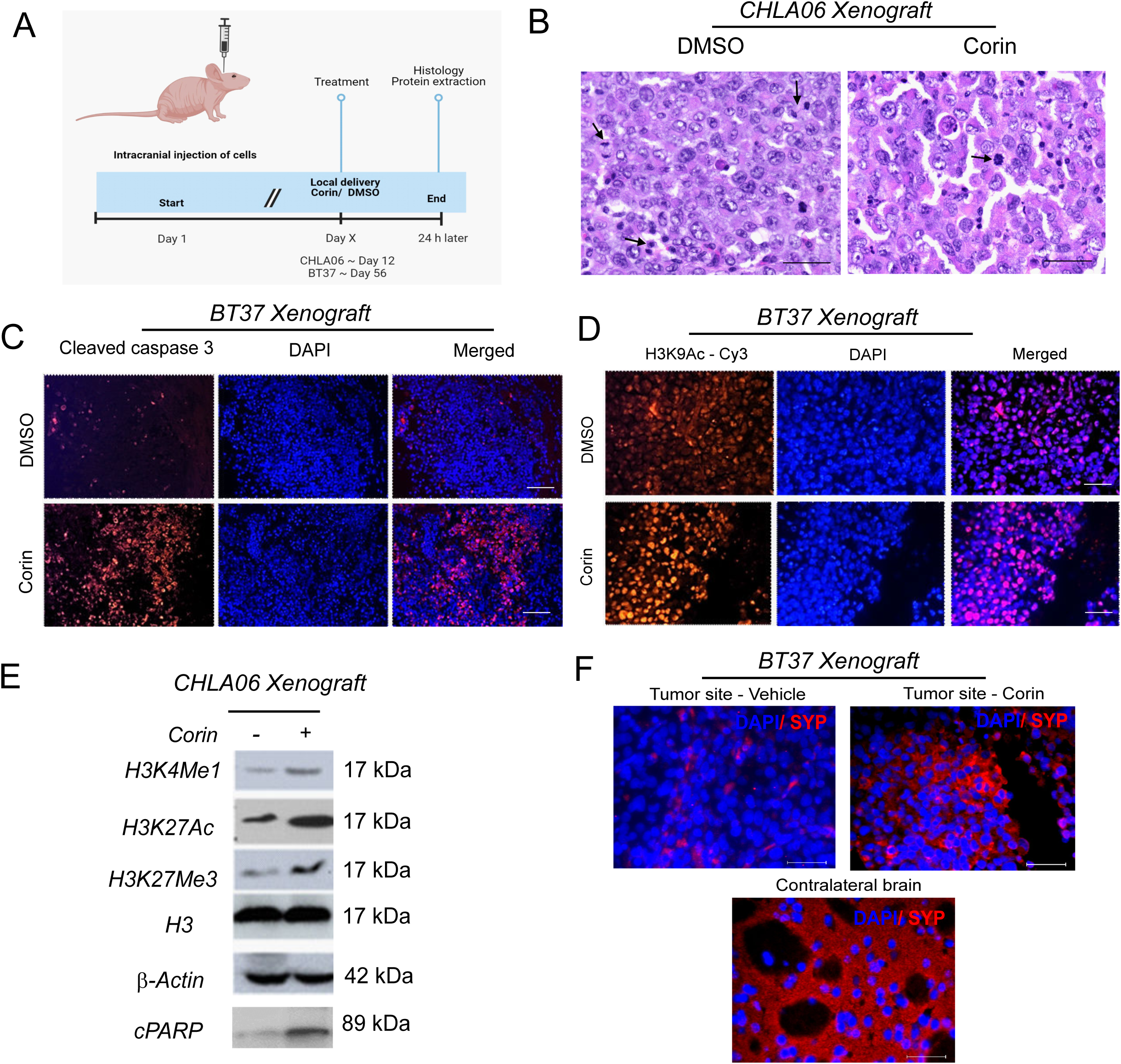
Short-term treatment of corin effectively hits the target and induces apoptosis in in vivo ATRT orthotopic xenograft models. **(A)** Schema showing the experimental design of short-term treatment of corin in mice. ATRT cells were injected through the guide-screw implanted in the right cerebral hemisphere and corin was injected through the guide screw after the mice started showing visible signs of tumor. Mice were sacrificed after 24 h and brains were extracted and processed for histology and Western blotting. [Created in BioRender. Geethadevi, A. (2024) https://BioRender.com/l53s982]. **(B)** Hematoxylin and eosin staining of mouse brains showing CHLA06 tumor xenografts in DMSO and corin-treated animals. Black closed arrows show mitotic figures. Scale 50 µm. **(C)** Immunofluorescence for cleaved caspase 3 (CC3) as a marker for apoptosis in BT37 orthotopic xenografts treated with DMSO or corin for 24 h. Scale 50 µm. **(D)** Immunofluorescence for H3K9Ac in BT37 orthotopic xenografts treated with DMSO or corin for 24 h. Scale 50 µm. **(E)** Western blot of H3K4Me1, H3K27Ac H3K27Me3, and cPARP in CHLA06 orthotopic xenograft tumors following 24 h DMSO or corin treatment. **(F)** Immunofluorescence for the differentiation marker, Synaptophysin, in BT37 orthotopic xenografts treated with DMSO or corin for 24 h. Contralateral brain is used as positive control. Scale 50 µm.

### Long-term treatment of corin decreased tumor cell growth and extended median survival in ATRT orthotopic xenograft models

To evaluate the effect of CoREST inhibition on ATRT tumor growth and survival in vivo, we established long-term orthotopic xenografts and treated them with corin (Figure 6A). We established intracranial orthotopic xenografts of BT37 in the right cerebral hemisphere, and we administered corin or vehicle intratumorally on a weekly basis using the guide screw method (see Supplementary Methods). To mimic the metastatic spread of ATRT, we injected CHLA06-ATRT cells in the lateral ventricle and then injected corin or vehicle into the intracerebroventricular (ICV) space weekly through the guide screw method (see Supplementary Methods). BT37 orthotopic xenografts treated with corin had significantly decreased tumor size with minimal to low leptomeningeal spread compared to the controls (Figure 6B, 6C). Corin similarly suppressed tumor growth of CHLA06 injected intraventricularly (Figure 6D, 6E). Weekly intratumoral corin extended the median survival of mice bearing BT37 orthotopic xenografts from 56 to 124 days (p = 0.007), with two mice surviving up to 152 days (Figure 6F). Furthermore, in the aggressive CHLA06-ATRT ICV xenografts, ICV treatment of corin led to an increase in median survival from 13 to 27 days with a significant increase in overall survival as measured by Log-rank test (p = 0.003) (Figure 6G). Corin was well-tolerated intratumorally as well as intracerebroventricularly, as there were no major changes in mouse body weights when administered long-term (Supplementary Figure 3A, 3B). In BT37 orthotopic tumor xenografts, we detected increased expression of H3K9Ac and H3K4Me1 5 days after corin treatment, demonstrating the persistent engagement of corin with histone targets (Supplementary Figure 3C, 3D). Consistent with our in vitro results, we found increased L1CAM and NEFL by immunofluorescence in corin-treated BT37 (Figure 6H, 6I) and CHLA06 orthotopic xenografts (Supplementary Figure 3E, 3F).

**Figure 6.**
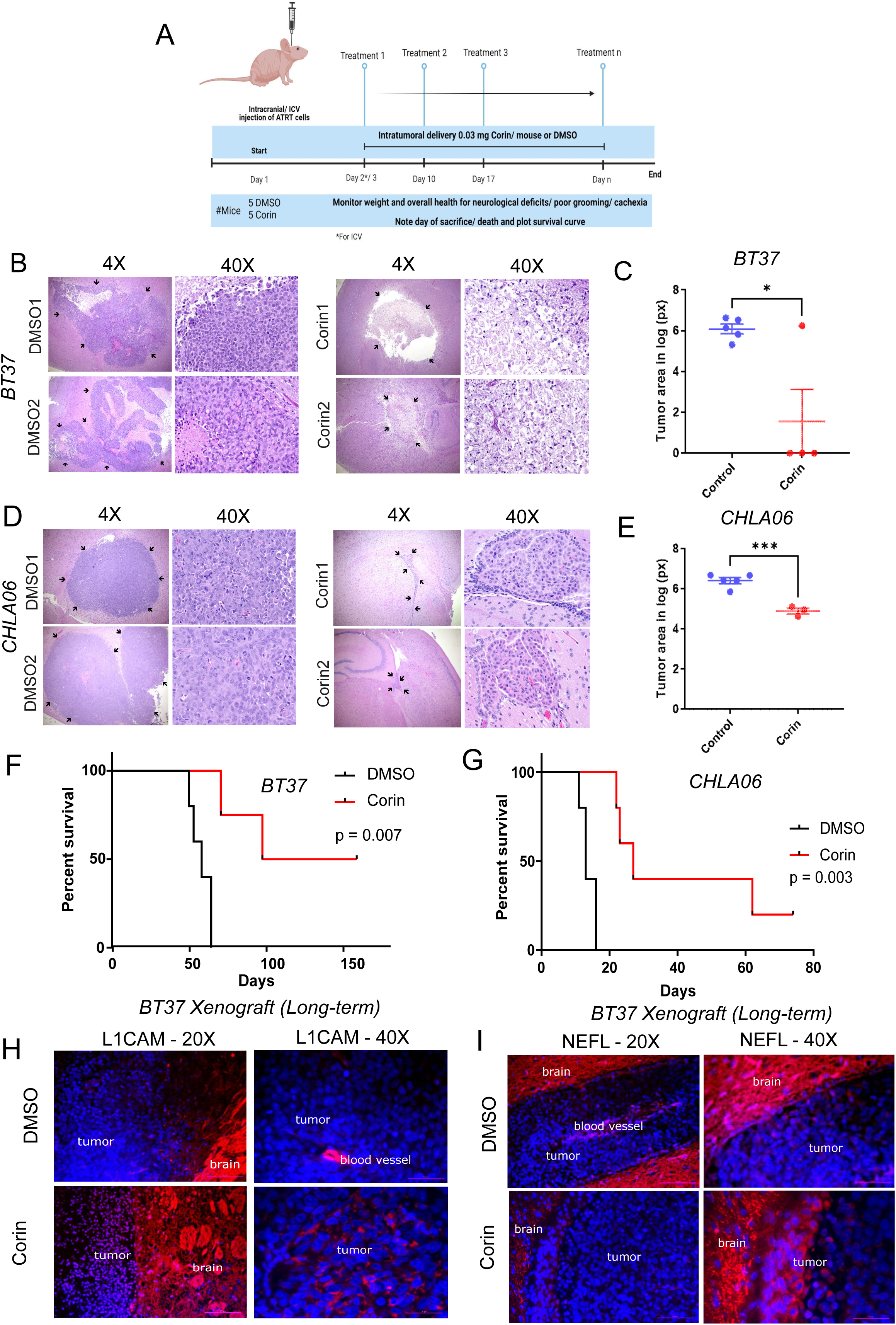
Long-term treatment of corin effectively extends the survival of mice bearing ATRT orthotopic xenograft tumors. **(A)** Schema showing the experimental design of long-term survival study. ATRT cells were injected through the guide-screw implanted in the right cerebral hemisphere (BT37) or through the ICV route (CHLA06) and DMSO or corin was injected through the guide screw after 2 days. Mice were treated with DMSO or corin every week and were sacrificed when they showed visible signs of tumor progression. Brains were extracted and processed for histology and immunofluorescence, and survival curves were plotted. [Created in BioRender. Geethadevi, A. (2024) https://BioRender.com/l53s982]. Hematoxylin and eosin staining and tumor measurements of mouse brains showing intracranial BT37 **(B, C)** and intracerebroventricular CHLA06 **(D, E)** tumor xenografts in DMSO and corin-treated animals following long-term treatment. Two representative animals are shown for each group. Graph of BT37 **(C)** and CHLA06 **(E)** tumor xenograft size as quantified using Qupath polygon annotation tumor xenografts in DMSO versus corin-treated animals following long-term treatment. **(F)** Graph of survival in mice with BT37 orthotopic xenografts treated with long-term administration of DMSO versus corin. Corin extended the median survival of mice bearing BT37 orthotopic xenograft from 56 to 124 days (p = 0.007). **(G)** Graph of survival in mice with CHLA06 intracerebroventricular tumor xenografts treated with long-term administration of DMSO versus corin. Corin extended the median survival of mice bearing CHLA06 orthotopic xenograft from 13 to 27 days (p = 0.003). Immunoflourescence for the differentiation markers, L1CAM **(H)** and NEFL **(I),** in BT37 orthotopic xenografts following long-term treatment with DMSO or corin. Statistical Analysis: unpaired, two-tailed student’s *t* test. All data represents mean ± SEM. * p ≤ 0.05, *** p ≤ 0.001.

## Discussion

ATRT is an epigenetically driven cancer, defined by loss of function of the SWI/SNF complex SMARCB1 protein, which results in reduced and aberrant SWI/SNF activity. While the SWI/SNF complex facilitates DNA accessibility, histone-modifying enzymes post-translationally alter the N-terminal tails of histones with consequent alterations in chromatin structure and binding of transcriptional regulatory proteins. The polycomb (PcB) group of repressors opposes SWI/SNF functions to create a balance between maintenance of a stem cell state and more differentiated neural precursors during development.^11^ The catalytic subunit of PRC2, EZH2, promotes methylation of lysine 27 on histone H3, a covalent chromatin modification that is associated with repressed heterochromatin and is increased in expression in SMARCB1-deficient cancers.^34^ EZH2 has been implicated as a critical driver of tumorigenesis in ATRT, and preclinical testing of the EZH2 inhibitor, tazemostat, demonstrated efficacy in 60% of rhabdoid tumor xenografts tested.^35^ However, clinical trials in pediatric ATRT patients with SMARCB1 mutations did not demonstrate significant clinical responses.^36^

Epigenetic therapy is an attractive tool for treatment of ATRT, however most epigenetic agents such as the pan HDAC inhibitor panobinostat are limited by their lack of specificity and limited therapeutic window.^37^ The CoREST complex inhibitor, corin, gains selectivity through its dual inhibitory activity versus LSD1 and HDAC1/2 and inhibited tumor growth to a greater extent that either an LSD1 or HDAC1/2 inhibitor alone or in combination.^15^

Recent studies have determined that critical gene loci occupied by the strong transcriptional repressor, REST (RE1 Silencing Transcription factor) are drivers of de-differentiation in ATRT.^12^ Furthermore, REST activity is largely mediated by association with the CoREST repressor complex.^15–17^ We therefore hypothesized that CoREST complex inhibition in ATRT would effectively reactivate critical REST-silenced differentiation leading to inhibition of tumor growth. Indeed, the IC_50_s for corin in our ATRT cell lines are less than those seen in melanoma or DMG.^18,21–23^ Weekly corin treatment of ATRT orthotopic xenografts in vivo suppressed ATRT tumor growth, without notable toxicity.

To improve corin delivery to the CNS, we used a guide-screw method that is often used for administration of oncolytic virus or immunotherapy in mouse orthotopic models.^38^ This method mimics the administration of drugs by an Ommaya reservoir, either directly to the tumor or to the CSF. Since most ATRT occur in the posterior fossa and most of the dissemination of the tumor is via the CSF, our establishment of the CHLA-06-ATRT model by intraventricular injection with subsequent suppression of growth by intraventricular administration of corin simulates how corin could be developed for clinical use in ATRT patients. Due to the high rate of relapse, many centers are now proposing maintenance therapy with intraventricular agents such as topotecan or liposomal cytarabine.^34,39,40^ Treatment with corin via an Ommaya reservoir may represent a novel epigenetic approach that allows treatment of patients with CSF dissemination at recurrence or maintenance therapy for patients who are at high risk of relapse.

In summary, corin demonstrates remarkable single agent activity against ATRT in orthotopic xenografts. By restoring epigenetic balance in ATRT, corin promoted differentiation and apoptosis and downregulated key stem cell factors that contribute to ATRT tumorigenesis. Moreover, corin upregulated pro-differentiation genes that are associated with favorable survival in human patients. These data suggest that corin may represent a novel treatment modality for patients with ATRT.

## Acknowledgments

RMA and MC are supported by Department of Defense Translational Research Award in Melanoma (W81XWH-21-1-0980); RMA and RJF are supported by Melanoma Research Alliance Grant 1045461; PAC is supported by NIH grant R35 GM149229. This project was supported by the Allegheny-Hopkins research fund, by the Giant Food Foundation, by the Alex’s Lemonade Stand Foundation and by the NCI core grant to Sidney Kimmel Comprehensive Cancer Center, Johns Hopkins University (EHR, JR, AG). We dedicate this manuscript to the memory of Francesca “Beans” Kaczynski, whose brave battle with ATRT inspired this collaborative effort.

## Author contributions

RMA conceived the project. AG, NV, MC, JR, EHR, and RMA designed the bench experiments. AG and EHR designed the animal studies. AG, NV, RJF, YD, and MC performed the bench experiments. AG and TF performed animal experiments. AG, NV, RJF, MC, KP, CGL, CGE, JX, JR, EHR, PAC and RMA analyzed and interpreted data. RJF, VK, and SB performed computational analyses. RMA, EHR, AG, NV, and MC wrote the manuscript. All authors discussed, revised, and approved the manuscript.

