## Supplementary Figure Legends for "The CoREST complex inhibitor, corin, leads to decreased tumor growth, increased cellular differentiation and extended lifespan in atypical teratoid rhabdoid tumor xenograft models"

**Supplementary Figure 1. Corin targets ATRT cells without affecting the iPSC-derived normal brain organoids. (A)** Graph and table showing the effective concentration of corin that induces cell death in ATRT cell lines in culture, as measured using the MUSE cell counter. **(B)** Graph and table showing the effective concentration of corin that reduces proliferation in ATRT cell lines in culture, as measured using the MUSE cell counter. **(C)** Corin reduces ATRT cell proliferation over 3-4 days compared to control as shown here by MTS growth assay. **(D)** Corin reduces PCNA expression in ATRT cell lines, BT37 and CHLA05, indicating reduced cell proliferation after corin treatment. Scale 50 µm. **(E)** Corin does not bear any toxic effect on iPSC-derived normal brain organoid as seen here in phase contrast microscopy (Scale 100 µm) as well as cPARP Western blot where cPARP is expressed in corin-treated CHLA06 but not in brain organoids.

**Supplementary Figure 2. Corin unwinds chromatin and differentiates ATRT cells along neuronal lineage. (A)** RNA-seq gene expression (Log2FoldChange) of genes with a "loss" or "gain" of ATAC-seq peaks in the gene body +/- 3Kbp of the gene in BT37 or CHLA06 cells treated with corin for 24 h compared to DMSO control. **(B)** Venn diagram showing the common upregulated genes between corin-treated BT37 and CHLA06. (**C**) K-means clustered heatmap of Z-scored DESeq2 normalized counts across each RNA-seq replicate. Each row represents a significant differentially expressed gene (DEG; $p < 0.05$, Log2FC $\geq 2$) from at least one cell line. Cluster 1 represents BT37-specific DEGs, cluster 2 represents CHLA06-specific DEGs, and cluster 3 represents common DEGs used for pathway enrichment analysis in Figure 3B. (D) GSEA analysis of DEGs in BT37 (left) and CHLA06 (right) using the Erkek (PMID: 30595504) gene set of SMARCB1 targets repressed in ATRT. **(E)** Immunoflourescence images showing MAP2 and Synaptophysin (SYP) in CHLA05 cells treated with corin or DMSO. **(F)** Immunofluorescence images showing Beta-3-tubulin in the ATRT cells treated with corin or DMSO.

**Supplementary Figure 3. Corin-administration removes repressive histone marks, induces neuronal differentiation and has no long-term side effects.** Long term administration of corin does not produce any observable side effects or toxicities as noted here by the mean body weights in **(A)** Intracranial BT37 ATRT xenograft, and **(B)** Intra-cerebroventricular CHLA06 ATRT xenograft. Immunofluorescence images showing H3K9Ac **(C)** and H3K4Me1 **(D)** expression in BT37 orthotopic xenografts collected from mice that died a week after corin treatment. Immunofluorescence staining showing the expression of L1CAM **(E)**, and NEFL **(F)** in CHLA06 orthotopic xenograft tumors after ICV treatment of corin.
