## Supplementary Methods for "The CoREST complex inhibitor, corin, leads to decreased tumor growth, increased cellular differentiation and extended lifespan in atypical teratoid rhabdoid tumor xenograft models"

### *Cell growth inhibition assay using MUSE cell counter*

The relative viability and the growth inhibition derived from the reduction in the number of viable cells after treatment with corin was evaluated using MUSE ViaCount Reagent, as per the manufacturer's instructions. ATRT cells were treated with corin (0.1 µM – 2 µM) or DMSO. After 72h, cells were re-suspended in their media inside the well and 20 µl of cell suspension was taken out into a MUSE eppendorf tube. 380 µl of ViaCount reagent was added to it, vortexed and the number of viable cells were counted using the MUSE cell counter. The percent viability and the total number of viable cells were both recorded. The relative percent viability and the relative number of viable cells were calculated taking the DMSO levels to be 100%. Plotting the relative cell viability data against the increasing doses of corin indicated the concentration of corin that induces half-maximal cell death while plotting the relative number of viable cell data against the increasing doses of corin yielded the concentration of corin that would inhibit the growth by 50%.

### *Annexin V/ Dead Cell assay using MUSE cell counter*

The proportion of cells undergoing various stages of apoptosis was estimated using Annexin V/ Dead cell kit using the manufacturer's instructions. ATRT cells were treated with corin (0.1 µM – 2 µM) or DMSO. After 72h, 100 µl of cell suspension was mixed with 100 µl of MUSE Annexin & Dead cell kit reagent and incubated for 20 min in dark. After setting appropriate gates to separate live from early and late apoptosis and dead cells in MUSE cell analyzer, apoptotic cells were counted in samples treated with and without corin. The number of cells undergoing various stages of apoptosis was recorded and relative values were calculated relative to DMSO.

***ATAC-Seq***

BT37 and CHLA06 cells were seeded 105,000 cells/$cm^2$ and treated with 1 µM Corin and DMSO for 24 h. 100,000 viable cells were aliquoted, and samples were prepared using ATAC-seq kit (Active Motif, 53150). Cells were pelleted at 500 x g for 5 min at 4 °C, washed with PBS and resuspended in ATAC Lysis Buffer followed by centrifugation at 500 x g 10 min at 4 °C. Pelleted nuclei were resuspended in Tagmentation Master Mix (Tagmentation Buffer, PBS, 0.01% Digitonin, 1% Tween 20, H2O and Assembled Transposomes) and incubated at 37 °C for 30 minutes in a thermomixer at 800 rpm. Then, DNA Purification Binding Buffer was added to the samples along with 3 M sodium acetate as required to reach the appropriate pH for binding. Samples were transferred to DNA purification columns, washed with wash Buffer, and eluted with DNA Purification Elution Buffer. Centrifugation of samples was done at 17000 x g for 1 min. Collected samples were amplified using a unique combination of index primers with the following cycling conditions: 72 °C for 5 min, 98 °C for 30s, 10 cycles of 98 °C for 10s, 63 °C for 30s and 72 °C for 1min, hold at 4°C. SPRI beads were added to each sample and vortexed, incubated for 5 min for binding, magnetically separated and washed with 80% ethanol solution twice while remaining attached. Following ethanol evaporation, beads were vortexed in DNA Purification Elution Buffer and incubated for 5 min at RT followed by collection of eluted samples.

### *ATAC-seq Analysis*

All reads were quality checked using FastQC v0.11.8 (Andrews 2010). Reads were aligned to the GRCh38 reference genome using bowtie2 v.2.4.2 (Langmead and Salzberg 2012). Bed and Bedgraph files were created using Bedtools v.2.31.0. Peaks were called using Macs2 v.2.2.7.1 with the nomodel, and nolambda options (Zhang et al. 2008). We used the AnnotatePeaks program present in hypergeometric optimization of motif enrichment (HOMER) tool (Heinz et al. 2010) to obtain the ATAC-seq signals for +/- 5kb around the Transcription start site (TSS) site. Bam files were normalized using RPKM to create BigWig files using the bamCoverage command in deeptools v.3.5.1. Profile plots were generated using the plotProfile function, and heatmaps were created using the plotHeatmap command. BigWig files were used to visualize individual gene tracks using IGV v.2.17.4. Genome annotation was completed using Samtools v.1.12 and homer v.4.1 with the command annotatePeaks.pl. BEDtools (Quinlan et al. 2010) and 'intersectBed' were used to identify the genes that gained/lost accessibility. Motif analysis was done using Samtools v.1.12 and homer v.4.1 using the findmotif.pl command searching for motifs within -400 to 100 bp of TSS.

### *RNA extraction*

RNA was extracted from ATRT cells treated with corin or DMSO using RNeasy Mini kit (Cat# 73404, Qiagen, Hilden, Germany) as per the manufacturer's instructions with some modifications. Briefly, cells were lysed using RNA lysis RLT buffer and passed through 1ml syringe attached to a 20 gauge needle at least 5 times. 70% ethanol was added and transferred into an RNeasy spin column. The RNA bound to the column was purified using RW and RPE buffers, and after a high-speed spin, finally collected in RNAse-free water. The quality and quantity of RNA extracted from the samples was measured using the Nanodrop 2000 (Thermofisher Scientific).

***RNA-seq Analysis***

RNA-seq was completed using Illumina NovoSeq 6000 platform. Low-quality reads and adaptors were trimmed using Trimmomatic v.0.36. Reads were quasi-mapped to the GRCh38 reference genome using Salmon v.1.1.0 (PMID: 28263959). Transcript counts were used for differential analysis using DESeq2. The Wald test was used to generate p-values and log2 fold changes. Genes with an adjusted p-value < 0.05 and absolute log2 fold change > 1.5 were called as differentially expressed genes for each comparison. Heatmap and hierarchical clustering for the RNA-seq was generated using ComplexHeatmap v.2.20.0 (Gu et al., 2016). Volcano plots were generated using ggplot2 and labeling was completed using ggrepel. GO Plots were generated using the enrichGO and dotplot commands in clusterProfiler v4.12.0 ( PMID: 22455463).

***Western immunoblotting***

30 µg of protein extracted from corin- or DMSO - treated ATRT cells or mouse xenografts were loaded onto 4-12% Bis-tris gel (Cat# NW04122F/ 20F, Thermo Fisher Scientific, Waltham, MA, USA) and separated by electrophoresis at 120 V. Proteins were transferred onto a methanol-activated PVDF membrane (Cat# 1620177, Bio-Rad Laboratories Inc.,

Hercules, CA, USA) at 2.5 A, 25V for 14 min using a fast transfer apparatus (Transblot Turbo Transfer System, Bio-Rad Laboratories Inc., Hercules, CA, USA). The membrane was then blocked in 5% BSA (Cat# 03116956001, Merck, NJ, USA), and incubated in primary antibody overnight followed by secondary antibody for 2 h at RT (see supplementary Tables 2 and 3 for antibody details). The blot was developed in ECL (Cat# NEL103001EA, Revvity Inc., Waltham, MA, USA; Cat# 34096, Thermo Fisher Scientific, Waltham, MA, USA) on X-ray films (Cat# BL1-810-100). The films were scanned and the optical density of the bands were quantified using Image-J software v1.54j. Beta actin was used as the normalizing control.

***Intracranial ATRT xenografts - Short-term corin treatment***

All the in-vivo experiments were conducted in athymic nude mice. Mice were anesthetized using 3:1 ratio of ketamine:xylazene in PBS. Injection guide holes were produced by an 18-gauge beveled needle in the right cerebral hemisphere and a guide screw was placed into these holes as described (Lal et al. 2000) using the following coordinates: antero-posterior = −3.5 mm; medio-lateral=2 mm; dorso-ventral = 3 mm (from the frontal suture). The guide was removed and BT37 ($1 \times 10^5$) and CHLA06 ($2.5 \times 10^4$) cells were injected in 3 µl of growth medium into the hole of the screw through a needle connected to a Hamilton syringe slowly so as not to induce sudden changes in cerebrospinal fluid pressure. The guide was plugged back into the hole, the skin was stapled securely and the mice were allowed to recover. After 3 days, the mice were randomly divided into two groups. The mice were monitored daily for their health, and when they showed visible signs of tumor such as neurologic deficits, poor grooming, and cachexia, corin (3 µl 0.03

mg/ mouse) or DMSO was administered through the guide screw. After 24h, the mice were sacrificed and their brains were dissected out. The tumor was located and a part of it was processed for Western blotting and the remaining for histology and immunohistochemistry/ immunofluorescence.

***Intracranial ATRT xenografts - Long-term survival study***

BT37 tumors were established in the right cerebral hemisphere using the guide-screw method as previously described. 3 days after the intracranial injection, the mice were randomly distributed into two groups, and corin or DMSO was administered weekly through the guide screw. The mice were monitored daily for signs of tumor growth and were sacrificed when they showed more than 20% loss of body weight and severe neurological deficits, and their brains were extracted for histology and immunohistochemistry or immunofluorescence. The date of their euthanasia was recorded and Kaplan Meier survival curve was plotted to assess the extension of median survival in mice bearing ATRT xenografts treated with corin.
