## Supplementary Tables for "The CoREST complex inhibitor, corin, leads to decreased tumor growth, increased cellular differentiation and extended lifespan in atypical teratoid rhabdoid tumor xenograft models"

**Supplementary Table 1. Cell lines and their culture conditions**

| Cell line | Culture type | Media | Composition of media |
|---|---|---|---|
| CHLA02-ATRT | Neurosphere (Suspension) | EF media | 70% DMEM (Cat# 11965-092), 30% Ham's F12 (Cat# 11765-054), 1% L-glutamine (Cat# 25030-081), 1% Antibiotic-Antimycotic (Cat# A5955, Sigma), 2% B27 supplement 50 X without Vit A (Cat# 12587-010), 0.25% Heparin (Cat# H3145-250KU), 0.02 µg/ml Epidermal Growth Factor/ EGF (Cat# AF-100-15-1MG, Peprotech), 0.02 µg/ml Fibroflast Growth Factor/ FGF (Cat# 100-18B) |
| CHLA04-ATRT | | | |
| CHLA05-ATRT | | | |
| CHLA06-ATRT | | | |
| CHLA266-ATRT | | | |
| BT12-ATRT | Adherent | RPMI media | 500 mL RPMI (Cat# 11875-093 ), 50 mL fetal bovine serum (FBS) (Cat# 16140-071), 1% L-glutamine (Cat# 25030-081), 1% Penicillin/Streptomycin (Cat# P0781, Sigma Aldrich) |
| BT37-ATRT | Semi-adherent | | |

**Supplementary Table 2. List of primary antibodies**

| Antibody name | Catalog No. | Antibody Dilution | |
|---|---|---|---|
| | | IF | WB |
| Anti-Beta 3 Tubulin | 5568[i] | 1:400 | - |
| Anti-Beta actin | sc-47778[ii] | - | 1:5000 |
| Anti-BrdU | 5292[i] | 1:400 | - |
| Anti-CC3 | 9661[i] | 1:400 | - |
| Anti-cPARP | 5625[i] | - | 1:1000 |
| Anti-H3K27Ac | 8173[i] | 1:500 | 1:1000 |
| Anti-H3K27Me3 | 9733[i] | 1:500 | 1:1000 |
| Anti-H3K4Me1 | 5326[i] | 1:500 | 1:1000 |
| Anti-H3K9Ac | 9649[i] | 1:500 | 1:1000 |
| Anti-L1CAM | 89861[i] | - | 1:1000 |
| Anti-LIN28A | 3978[i] | - | 1:1000 |
| Anti-LIN28B | 4196[i] | - | 1:1000 |
| Anti-MAP2 | 8707[i] | 1:400 | - |
| Anti-NEFL | 2837[i] | - | 1:1000 |
| Anti-PCNA | 10205-2-AP[iii] | 1:400 | - |
| Anti-SOX2 | MA1-014[iv] | - | 1:1000 |
| Anti-SYP | 17785-1-AP[iii] | 1:400 | - |
| Anti-Total H3 | 4499/ 14269[i] | - | 1:1000 |

[i]Cell Signaling Technologies, MA, USA; [ii]Santacruz Biotechnologies, TX, USA; [iii]Proteintech Group Inc., IL, USA; [iv]Sigma Aldrich, MO, USA

**Supplementary Table 3. List of secondary antibodies**

| Antibody name | Catalog No. | Antibody Dilution |
|---|---|---|
| **Western Immunoblotting** | | |
| Anti-rabbit | 7076[i] | 1:3500 |
| Anti-mouse | 7074[i] | 1:5000 |
| **Immunofluorescence** | | |
| Anti-rabbit-Cy3 | 711-165-152[ii] | 1:500 |
| Anti-mouse-Cy3 | 715-165-151[ii] | 1:500 |

[i]Cell Signaling Technologies, MA, USA; [ii]Jackson Immunoresearch Laboratories Inc., Westgrove, PA, USA

**Supplementary Table 4. ChIP-qPCR antibodies and primers**

| Antibody name | Catalog No. | Antibody Dilution |
|---|---|---|
| IgG | 2729[i] | 10 µg/sample |
| LSD1 | ab17721[ii] | 10 µg/sample |
| RCOR1 | 07-455[iii] | 10 µg/sample |
| H3K9ac | ab32129[ii] | 4 µg/sample |

[i]Cell Signaling Technologies, MA, USA; [ii]Abcam, UK; [iii] EMD Millipore, MA, USA

| Primer Target | Forward (5’ -> 3’) | Reverse (5’ -> 3’) |
|---|---|---|
| *NEFL* | GCCGTTCTGCCACCCCTATT | CGGCGTGCCGTGATCG |
| *L1CAM* | CCGCTGTGAAAGCTCGGA | TTCCTTGCCGGACTCCTCTC |

**Supplementary Table 5. Dose of corin used for ATRT cell lines in each experiment**

| **Experiment name; <u>Time point</u>**<br>***Cell Line*** | **Dose of corin** |
|---|---|
| **Western Immunoblotting of cPARP; <u>72h</u>** | |
| *BT37* | 1 µM |
| *CHLA05* | 500 nM |
| *CHLA06* | 1 µM |
| *iPSC-derived brain organoid* | 5 µM (Tested 1-10 µM) |
| **Western Immunoblotting of histone markers; <u>24h</u>** | |
| *BT37* | 1 µM |
| *CHLA05* | 500 nM |
| *CHLA06* | 1 µM |
| **BrdU immunoassay; <u>72h</u>** | |
| *BT37* | 500 nM |
| *CHLA05* | 250 nM |
| *CHLA06* | 1 µM |
| **CC3 immunostaining; <u>72h</u>** | |
| *BT37* | 500 nM |
| *CHLA05* | 250 nM |
| *CHLA06* | 1 µM |
| **MUSE Annexin V assay; <u>24h</u>** | |
| *BT37* | 1 µM |
| *CHLA05* | 500 nM |
| *CHLA06* | 1 µM |
| **ATAC-seq; <u>24h</u>** | |
| *BT37* | 1 µM |
| *CHLA06* | 1 µM |
| **RNA-seq; <u>24h</u>** | |
| *BT37* | 1 µM |

| | |
|---|---|
| *CHLA06* | 1 µM |
| **WGA Staining, Immunostaining for neuronal differentiation factors** | |
| *BT37* | 1 µM |
| *CHLA05* | 500 nM |
| *CHLA06* | 1 µM |
| **Western immunoblotting for SOX2, LIN28A, NEFL; 72h** | |
| *BT37* | 1 µM |
| *CHLA05* | 500 nM |
| *CHLA06* | 1 µM |
| **Western immunoblotting for LIN28B, L1CAM; 72h** | |
| *BT37* | 1 µM |
| *CHLA05* | 500 nM |
| *CHLA06* | 1 µM |
| **MTS Assay; 0 – 96/ 120h** | |
| *BT37* | 500 nM |
| *CHLA05* | 250 nM |
| *CHLA06* | 1 µM |
| **PCNA immunostaining; 72h** | |
| *BT37* | 500 nM |
| *CHLA05* | 250 nM |
| *CHLA06* | 1 µM |
