## Supplementary Figures for "The CoREST complex inhibitor, corin, leads to decreased tumor growth, increased cellular differentiation and extended lifespan in atypical teratoid rhabdoid tumor xenograft models"

### Supplementary Figure 1

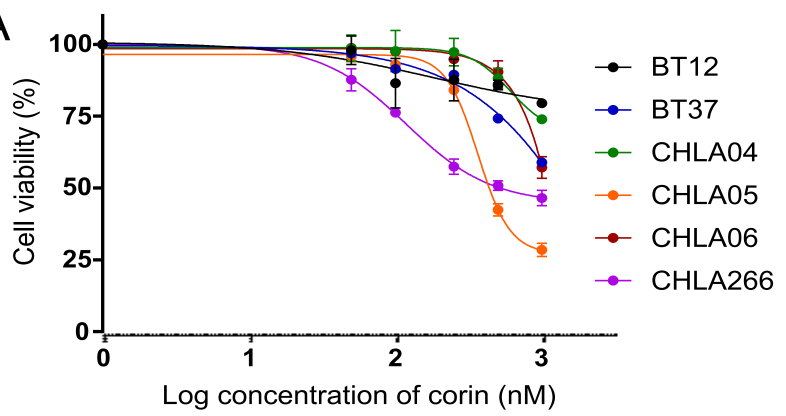


| *ATRT Cell line* | *IC50 of Corin (μM)** |
|---|---|
| BT37 | 0.902 |
| CHLA04 | 1.2 |
| CHLA05 | 0.330 |
| CHLA06 | 1.327 |
| CHLA266 | 0.334 |
| BT12 | 2.5 |

*Concentration of Corin that reduces cell viability by 50%. Calculated based on viable cells relative to DMSO using MUSE ViaCount assay.

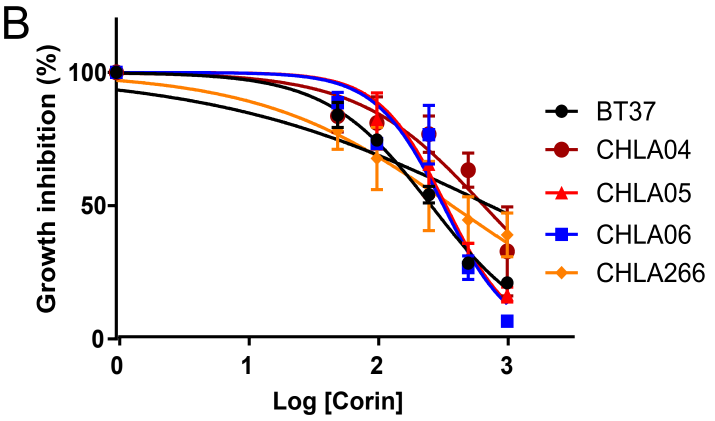


| *ATRT Cell line* | *IC50 of Corin (μM)** |
|---|---|
| BT37 | 0.261 |
| CHLA04 | 0.661 |
| CHLA05 | 0.339 |
| CHLA06 | 0.323 |
| CHLA266 | 0.377 |

*Represents the concentration of Corin that inhibits cell growth by 50%, and is calculated based on total viable c cell count after 72 h.

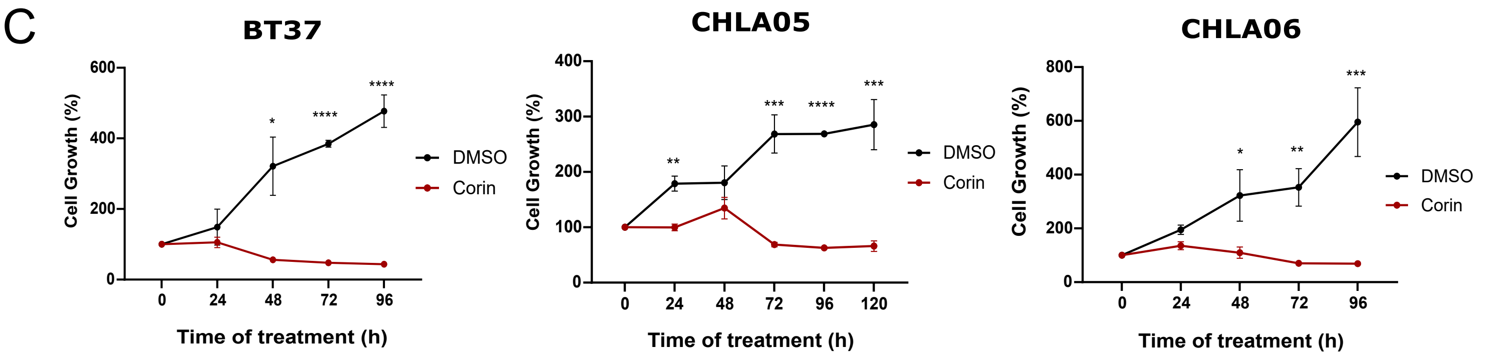


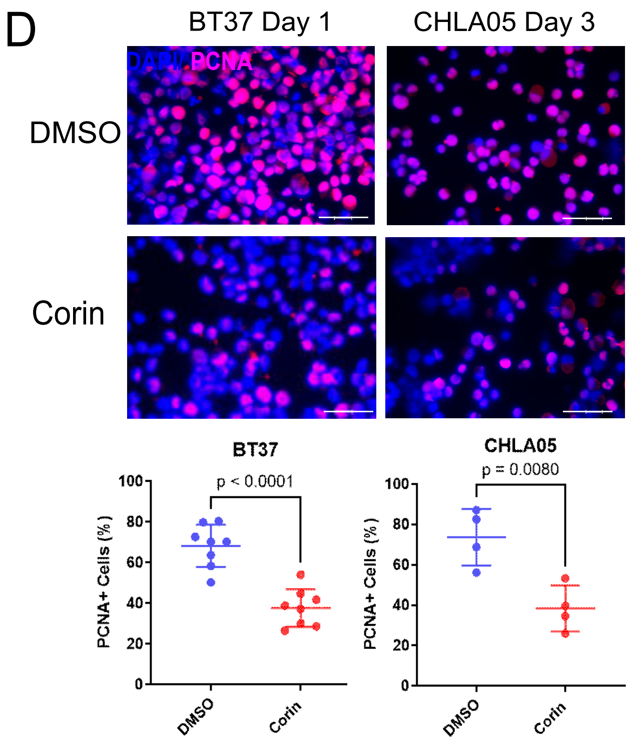


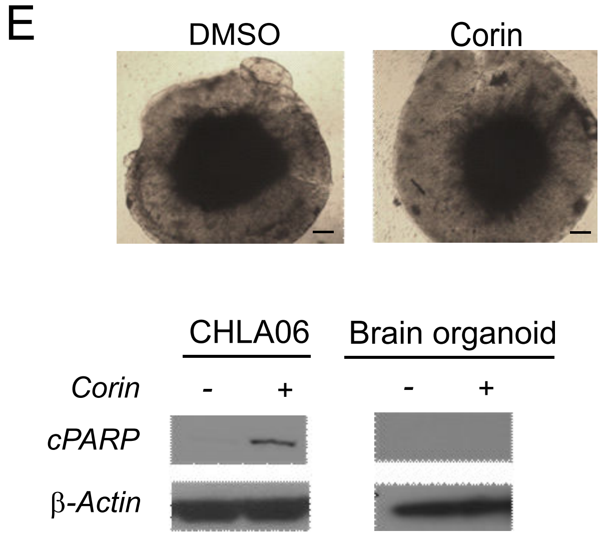

#### Supplementary Figure 2

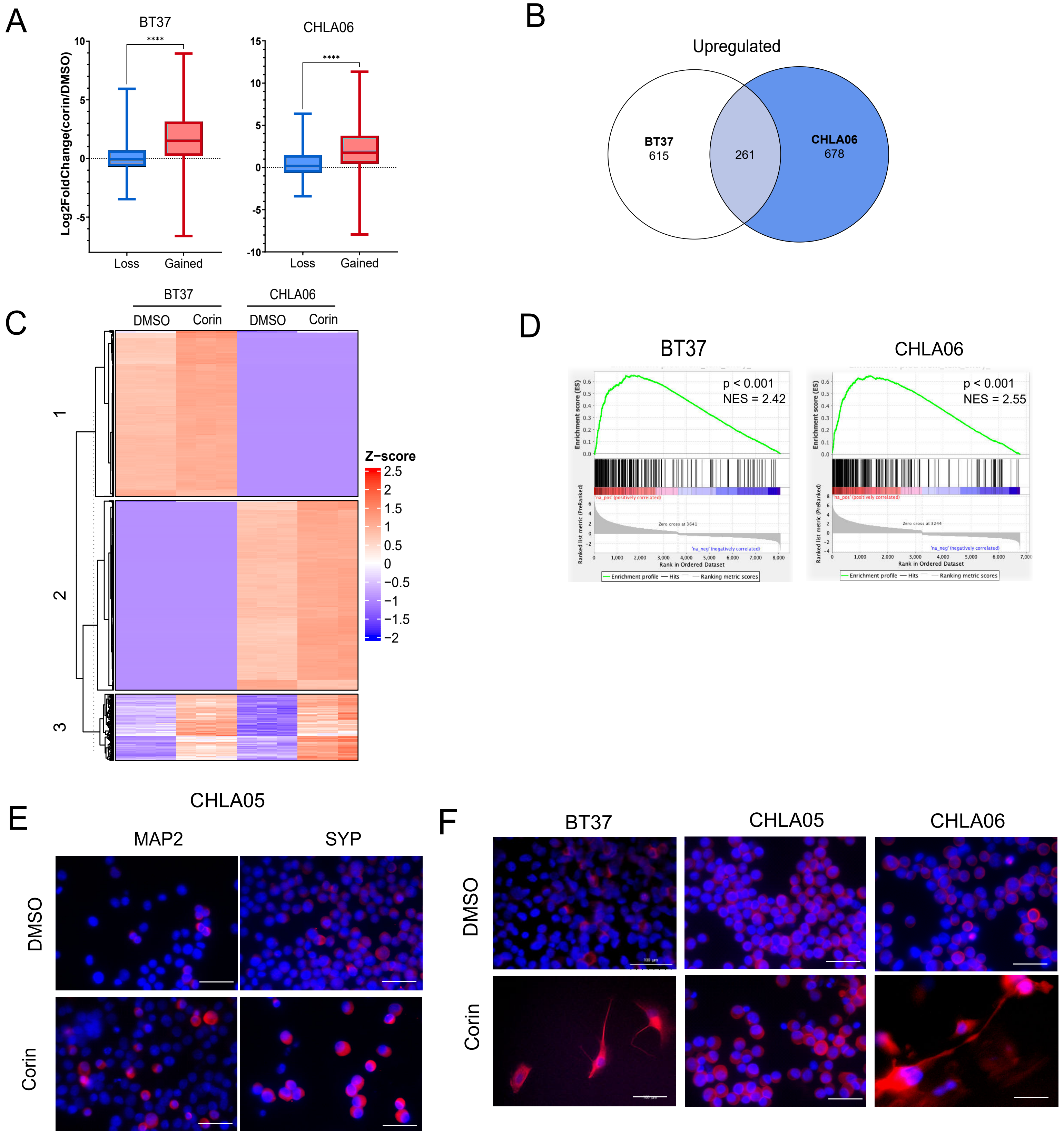

**Supplementary Figure 3**

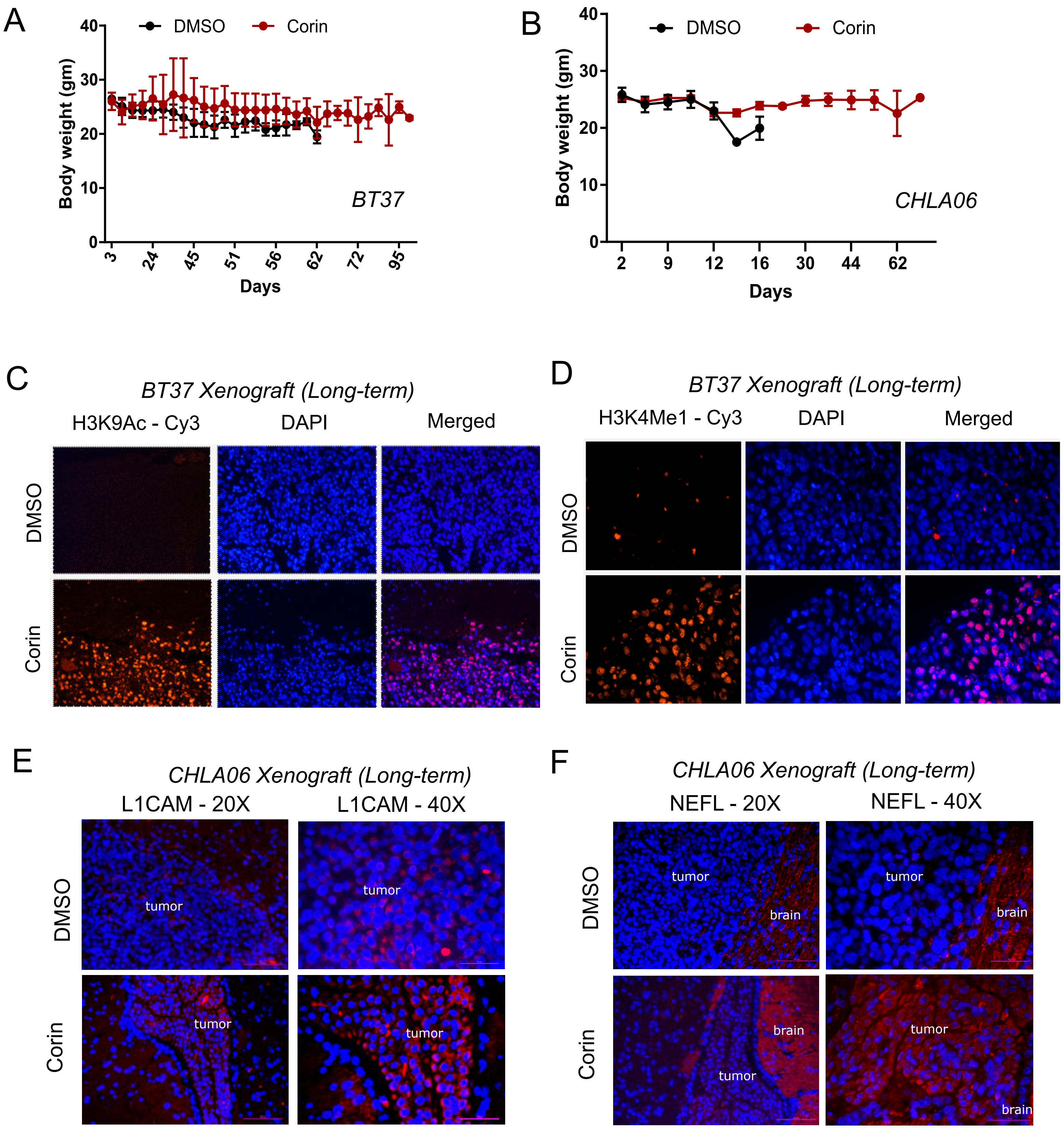
